# Optic nerve crush does not induce retinal ganglion cell loss in the contralateral eye

**DOI:** 10.1101/2024.09.16.613181

**Authors:** Florianne E. Schoot Uiterkamp, Margaret E. Maes, MohammadAmin Alamalhoda, Arsalan Firoozi, Gloria Colombo, Sandra Siegert

## Abstract

**Purpose:** Optic nerve crush (ONC) is a model for studying optic nerve trauma. Unilateral ONC induces massive retinal ganglion cell (RGC) degeneration in the affected eye, leading to vision loss within a month. A common assumption has been that the non-injured contralateral eye is unaffected due to the minimal anatomical decussation of the RGC projections at the chiasm. Yet, recently, microglia, the brain-resident macrophages, have shown a responsive phenotype in the contralateral eye after ONC. Whether RGC loss accompanies this phenotype is still controversial.

**Methods:** Using the available RGCode algorithm and developing our own RGC-Quant deep-learning-based tool, we quantify RGC’s total number and density across the entire retina after ONC.

**Results:** We confirm a short-term microglia response in the contralateral eye after ONC, but this did not affect microglia number. Furthermore, we cannot confirm the previously reported RGC loss between naïve and contralateral retinas five weeks after ONC induction across the commonly used Cx3cr1^creERT2^ and C57BL6/J mouse models. Neither sex nor the direct comparison of the RGC markers Brn3a and RBPMS, with Brn3a co-labeling, on average, 89% of the RBPMS^+^-cells, explained this discrepancy, suggesting that the early microglia-responsive phenotype does not have immediate consequences on the RGC number.

**Conclusions:** Our results corroborate that unilateral optic nerve injury elicits a microglial response in the uninjured contralateral eye but without RGC loss. Therefore, the contralateral eye should be treated separately and not as an ONC control.

## Introduction

Retinal ganglion cells (RGC) relay the detected and processed light information within the retina for further visual processing in the brain. Their axons bundle at the optic disc, forming the optic nerve, which extends to the eye’s contralateral brain areas and decussates only minimally at the chiasm in mice ^1^. Blunt force, penetrative trauma, or an increase in ocular pressure can induce an injury to the optic nerve, causing RGC axon degeneration and vision loss ^2^. A standard model to mimic this optic nerve trauma is the optic nerve crush (ONC) ^3,4^, where the exposed optic nerve is pinched with a forceps. This procedure results in approximately 80% RGC death within 14 days in the ONC-receiving eye and elicits a pronounced glial response ^5,6^. During this phase, microglia adopt a responsive phenotype and promote the release of pro-inflammatory cytokines and chemokines ^7–9^. Frequently, the non-injured, contralateral eye has been used as an internal control under the assumption that due to the low decussation between both eyes at the chiasm, the effects in the contralateral eye will be limited ^7,10–15^. However, recent studies suggested that unilateral ONC induces a mild responsive phenotype in microglia, increases their proliferation, and upregulates activation-specific proteins and inflammatory markers in the contralateral eye ^3,16–18^.

Besides the microglia phenotype, several studies analyzed potential consequences on the RGC number, but the authors did not find any changes even six months after ONC ^19–21^. This is in contrast to other reports, which found up to 20% RGC decrease three weeks after ONC ^22,23^. Specifically, Lucas-Ruiz et al. (2019) found a 15% RGC loss and suggested a connection to the inflammation signature. They rescued the RGC loss only in the contralateral eye upon systemic treatment with either the tetracycline antibiotic minocycline, which selectively targets microglia or the non-steroidal anti-inflammatory drug meloxicam.

Inflammation has been shown to influence human eye surgical procedures. The chance of corneal transplant rejections increases threefold if the patient has already undergone the same procedure previously in the opposite eye ^25,26^. In light of a potential concern that an injury in one eye might severely impact the vision in the other, a special warrant has been made to clarify this aspect. Thus, we decided to investigate commonly used RGC antibodies and quantify the RGC number and density across the retina in a minimally biased approach using automized deep-learning-based tools. We confirmed the microglia-responsive phenotype, yet we did not observe RGC loss in the contralateral retina after ONC investigating two mouse models, sex, and RGC-selective markers, RBPMS and Brn3a. Thus, unilateral injury to the optic nerve results in glial activation in the contralateral eye, which does not lead to neuronal loss.

## Materials and Methods

### Animals

All animal housing and procedures were approved by the “Bundesministerium für Wissenschaft, Forschung und Wirtschaft (bmwfw) Tierversuchsgesetz 2012, BGBI. I Nr. 114/2012 (TVG 2012)” under GZ: 2021-0.607.460. If not otherwise indicated, adult mice (3-5 months) of both sexes were used. Mouse strains C57Bl6/J (#000664), *Cx3cr1*^CreERT2^ (#020940), and PhAM^fl/fl^ (#018385) were obtained from Jackson Laboratories. For FACS sorting, PhAM^fl/fl^ mice containing a mitochondrial-specific green fluorescent tag were crossed with *Cx3cr1*^CreERT2^ mice to ensure the microglia specificity of the tag ^27^. Since *Cx3cr1*^CreERT2^ is a knock-in/ knock-out model, the *Cx3cr1*^CreERT2^ and *Cx3cr1*^CreERT2^/ PhAM^fl/fl^ mouse lines were always used heterozygous crossed with C57Bl6/J. Mice were housed in the ISTA preclinical facility in a 12-hour light-dark cycle in individually ventilated cages in a controlled environment with food and water provided *ad libitum*.

### Tamoxifen administration for Cre-induced recombination

30 mg/mL Tamoxifen (Sigma Aldrich, T5648-5G) was dissolved in 90% corn oil (Sigma Aldrich, C8267-500ML) and 10 % ethanol and sonicated for 45 minutes. Adult Cx3Cr1^creERT2^ and *Cx3cr1*^CreERT2^/ ^PhAMfl/fl^ heterozygous mice were injected intraperitoneally daily (150mg/kg) for three consecutive days to induce Cre activity. Cx3CR1 is also expressed by circulating monocytes, peripheral macrophages, and dendritic cells ^28^. Naïve and ONC mice were injected with tamoxifen due to the observation of some tamoxifen-independent activity of Cre in reporter lines ^29^. To ensure that the expression of creERT2 is limited to microglia, experiments were performed at least three weeks after tamoxifen induction, using the turnover rate of peripheral cells ^30,31^.

### Optic nerve crush (ONC) procedure

During surgery, the mice were anesthetized with 2.5% (v/v) isoflurane (Zoetis) supplied with oxygen at a flow rate of 0.6 L/min, and their body heat was maintained at 37°C using a heating pad. To numb the eye, Proparacaine hydrochloride 0.5% eye drops (Ursapharm Arzneimittel GmbH) was applied, and to alleviate the pain, a subcutaneous injection of 5mg/kg Metacam (Meloxacam, Boehringer Ingelheim) or 0.1 mg/kg Buprenorphine (Bupaq, Richter Pharma) was given. The optic nerve was exposed intraorbitally and pinched using a curved N7 self-closing forceps (Dumont) for 10 s, approximately 1 mm from the posterior pole. Eye ointment was applied post-operatively to protect the cornea from dehydration (OLEOvital). The injured ONC eye was used to verify the sufficiency of the crush on day five, as published previously ^27^.

### Perfusion and Tissue Processing

Animals were anesthetized with isoflurane (Zoetis) and transcardiacally perfused with 20 mL of phosphate-buffered saline (PBS) with heparin (100 mg/L, Sigma H0878), followed by 20 ml of 4% (w/v) paraformaldehyde (PFA, Sigma P6148) in PBS using a peristaltic pump (Behr PLP 380, speed: 26 rpm). The animals were decapitated, and the eyeballs with optic nerves were removed and post-fixed in 4% (w/v) PFA/PBS for 30 minutes. Then, the tissue was washed 3x in PBS and stored at 4°C. Retinas were dissected from the enucleated eyes in PBS and then transferred to 30% (w/v) sucrose (Sigma 84097) in PBS overnight at 4°C.

### Immunohistochemistry

Retinas underwent three freeze/thaw cycles on dry ice to increase antibody permeability. Next, a blocking step was performed using a blocking solution containing 1% (w/v) bovine serum albumin (Sigma A9418), 5% (v/v) Triton X-100 (Sigma T8787), 0.5% (w/v) sodium azide (VWR 786-299), and 10% (v/v) donkey serum (Millipore S30) for 1 hour at room temperature on a shaker. Primary antibodies were diluted in an antibody solution containing 1% (w/v) bovine serum albumin, 5% (v/v) triton X-100, 0.5% (v/v) sodium azide, 3% (v/v) donkey serum, and incubated for two nights on a shaker at 4°C. The following primary antibodies were used: rat α-CD68 (cluster of differentiation 68, AbD Serotec MCA1957, clone FA-11, Lot 1807, 1:250); goat α-Iba1 (ionized calcium-binding adapter molecule 1, AIF1, Abcam, ab5076, Lot FR3288145-1, 1:250); rabbit α-Iba1 (GeneTex, GTX100042, Lot 41556, 1:750); mouse α-Brn-3a (Brain-specific homeobox/POU domain protein 3A, Merck, MAB1585, Lot 3942133, 1:100); guinea pig α-RBPMS (RNA Binding Protein with Multiple Splicing, PhosphoSolutions, 1832-RBPMS, Lot NB322g, 1:200); rabbit α-RBPMS (Abcam, ab194213, Lot GR3383144-6, 1:200). Following three washes with PBS, the samples were incubated protected from light for 2 hours at room temperature on a shaker with the secondary antibodies diluted in antibody solution. The secondary antibodies raised in donkeys were purchased from Thermo Fisher Scientific (Alexa Fluor 488, DyLight 650, Alexa Fluor 568, Alexa Fluor 647, 1:2000) and Jackson ImmunoResearch (Alexa Fluor 647, 1:1000). After three washes with PBS, the samples were incubated with Hoechst 33342 (Thermo Fisher Scientific, H3570, 1:5000) diluted in PBS for 30 minutes. The retinas were flat mounted on microscope glass slides (Assistant, 42406020) with coverslips (Menzel-Glaser #0) using an antifade solution (10% (v/v) Mowiol (Sigma, 81381), 26% (v/v) glycerol (Sigma, G7757), 0.2M tris buffer pH 8, 2.5% (w/v) Dabco (Sigma, D27802)). Parafilm spacers were put between the glass slides and coverslips to prevent tissue distortion. For immunolabeling the RGCs, the retinas were first flat mounted on Superfrost plus adhesion microscope slides (Epredia, 12625336). After flat mounting, retinas were post-fixed again for 1 hour to flatten them and then further processed for immunohistochemistry as described above.

### Confocal microscopy

#### Microglia analysis

Images for CD68 quantification, microglia density, and Sholl analysis were acquired with a Zeiss LSM800 upright or Zeiss LSM900 upright using a Plan-Apochromat 40x oil immersion objective NA 1.3. Tiled (2×2) images with a resolution of 0.156×0.156×0.3 µm were acquired from two opposing quadrants of the retinas to correct for any intrinsic variability of microglial response to injury. Inner plexiform layer microglia were selected by setting the z-stack between the ganglion cell and inner nuclear layer Hoechst staining.

#### Retinal ganglion cell (RGC) analysis

Images for RGC quantification were acquired on an Andor Dragonfly 505 spinning disk using a CFI P-Apo 20×objective NA 0.75 and 25 µm pinhole size with a resolution of 0.603×0.603×0.494 µm. The size of the z-stack tile image was set up to capture the entire flat-mounted retinas.

### Image analysis

Confocal images were converted to .ims files using the Imaris converter and stitched using the Imaris stitcher. Unless otherwise specified, the analysis was performed with IMARIS version 9.1-9.3 (Bitplane).

#### CD68 volume within microglia

Three-dimensional surfaces were generated for microglia and CD68 using the surface rendering module of Imaris 9.3 with the surface detail set to 0.2 µm. The *surface-surface coloc* plugin was used to obtain the CD68 surface within microglia. The total percentage of CD68 volume within this microglia volume was calculated per quadrant and represented as the mean of two retinal quadrants.

#### Microglia density

Iba1^+^-microglia were counted using Imaris’s *spot-detection function*. The total cell count was normalized to the entire image area and represented as the number of cells per mm^2^.

#### Filament tracing and Sholl analysis

Microglia morphology was traced in three dimensions using the *Filament* Wizard in Imaris. The starting point was detected with a diameter of 10 µm, seeding points were set to 0.6 µm, and the disconnected segment was removed. Final filament traces were manually edited to remove incorrect segments from the semi-automatic *Filament* tracing. The ‘Number of Filament Sholl intersections’ from the *Statistics* file was used to plot the number of intersections per increasing radii of 1 µm. Plotted Sholl curves were truncated at 60 µm.

### Automated RGC counting

#### Counting RBPMS^+^-cells using RGCode

Stitched .ims files were transferred to Fiji (https://imagej.net/software/fiji). Channels were split, and the RBPMS channel was selected. Then, the Z-stack was converted to a maximum-intensity Z-project. The image background was subtracted with a rolling ball radius of 50 px, and median-filtered with a radius of 1 px, and the local contrast was enhanced. The image was saved as a .tif file. The RBPMS^+^-RGC were counted using the RGCode script as described ^32^. RGCode provides the total count, the cell density, and the retinal area as output.

#### Counting Brn3a^+^-cells using RGC-Quant

The model was trained using datasets labeled semi-automatically using Imaris. The datasets comprised 3D confocal images of RGCs, which were segmented into 3D cubes for training purposes.

#### Training process

*1. Training data gathering*. The training datasets were gathered from six retina samples in which RGCs were semi-automatically labeled using Imaris. We have used an accumulated number of ‘245,000’ labeled RGCs for the training procedure. *2. Data decomposition*: Because of the input size limitation in the 3D U-Net model, we decomposed each confocal image into smaller cubes to serve as model input. This decomposition ensured that all parts of the original images could be reconstructed from these cubes. *3. Training procedure*: The 3D U-Net model was trained on these cubes using a supervised learning approach. The loss function combines dice loss and binary cross-entropy, which helps handle class imbalance and improves segmentation accuracy. *4. Validation*: A separate validation set, comprising 40% of the total dataset, was used to monitor the model’s performance during training and to fine-tune hyperparameters. Early stopping and model checkpointing were implemented to prevent overfitting. *5. Performance evaluation*: To measure the performance of the RGC counting model, we employed several metrics, including precision, recall, F1-score, and the Dice coefficient. These metrics were computed by comparing the model’s predictions against semi-automatic annotated ground truth data. Additionally, the counting accuracy was evaluated by comparing the automatic counts with manual counts performed by expert annotators. The results showed a high correlation between automatic and manual counts, demonstrating the model’s reliability and accuracy. After training, the model was applied to new datasets for detecting Brn3a^+^-cells.

#### Detection process

*1. Cell detection*: The trained 3D U-Net model was used to detect cells within the 3D segments of new confocal images. *2. Post-Processing*: Post-processing techniques, such as morphological operations, were applied to refine the detected cell regions and exclude non-cellular structures like blood vessels and fibers. *3. Counting Cells*: The number of cells was determined by counting the number of connected components in the segmented images.

This combined approach enabled us to detect cells and automatically compute cell numbers, ensuring high precision and reliability in quantifying RGCs from confocal images.

### Reverse transcription-quantitative real-time PCR (RT-qPCR) and gene expression analysis

Real-time PCR was performed as described previously ^27^. Retinal microglia from *Cx3cr1*^CreERT2/C57BL6/J^/PhAM^fl/fl^ mice were FACSorted using a SONY SH800SFP. From each retina, 100 microglia were sorted and cDNA synthesized with the NEBNext® Single Cell/Low Input cDNA Synthesis & Amplification Module (New England Bio Labs, #E6421L) according to the manufacturer’s protocol. Primers for CD68 (FW: 5’-CCTCTGTTCCTTGGGCTATAAG-3’, REV: 5’-ATTGAGGAAGGAACTGGTGTAG-3’) and GADPH (FW: 5’-ACAGCAACTCCCACTCTTC-3’, REV: 5’-CATTGTCATACCAGGAAATGAGC-3’) were validated based on their efficiencies. For gene expression analysis, RT-qPCR (Luna® Universal qPCR Master Mix; New England Biolabs; M3003L) was performed in 384-well plates (Bio-Rad; HSR4805) on a Roche Lightcycler 480 according to the manufacturer’s manual. PCR reactions were carried out in triplicate, from which the mean Cq value was calculated. Fold change differences between the ONC and the naïve conditions were calculated using the delta-delta Ct method ^33^. dCq values were obtained by normalizing mean Cq values to the reference housekeeping gene (GAPDH) measured within the same experiment (Equation 1). ddCq values were then calculated by normalizing dCq values to the control condition (Equation 2). Fold changes were obtained by transforming ddCq values from log2-scale to linear scale (Equation 3). Fold changes were used for data visualization.

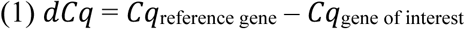

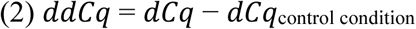

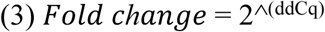

### Statistical analysis

All statistics were performed using R (version 3.4.4), as indicated in the figure legends. Groups for comparison were first tested for normal distribution and equal variance using the Shapiro-Wilk test and Levene test, respectively. Where applicable, post hoc tests were performed using Tukey’s multiple comparisons of means or the Conover-Iman test for non-parametric multiple comparisons. For Sholl analysis comparisons, linear regression models were performed using the *lme4* package. The default contrast for unordered variables was changed to ‘contr.sum’, allowing type III ANOVA comparison. Descriptive statistics were generated using the *psych* package. Data were presented as the mean ± SD. Plots were generated using ggplot2 (version 3.5.1). Error bars represent the standard error of the mean. Significance levels are indicated using the following notation (ns: non-significant with *p* > 0.05; *\* p* < 0.05; ** *p* < 0.01; *\*\*\* p* < 0.001).

## Results

### Microglia in the contralateral eye after optic nerve crush

Multiple studies have investigated the microglia phenotype in the contralateral eye after optic nerve crush (ONC) and confirmed their responsive phenotype across different mouse or rat strains such as C57Bl6/J, BALB/cJ, and Sprague-Dawley or Brown-Norway rats ^21–24,34,35^. To assess the microglia phenotype, we used transgenic mice strains *Cx3cr1*^creERT2/C57BL6/J^, which harbors a tamoxifen-inducible CRE recombinase in the macrophage-specific *Cx3cr1* locus. This mouse model is widely used to study microglia because it allows the crossing with reporter mouse lines for either distinct fluorophore visualization or selective gene knockout, or it increases transduction efficiency with a *CRE*-dependent virus ^27,30,36,37^. We performed a unilateral ONC injury (**Figure 1A**) and sacrificed the mice 2, 5, 7, 14, or 35 days after ONC. For the non-injured contralateral eye, we performed immunostaining for Iba1 and the endosomal-lysosomal marker CD68. CD68 is commonly upregulated in microglia upon a stress response and indicates a responsive microglia phenotype ^38,39^. We imaged microglia in the inner plexiform layer (IPL) due to their proximity to the RGC and quantified the CD68 volume within microglia. The CD68 level significantly increased within the first week after ONC and gradually declined to the level of the naïve animal at 35 days (**Figure 1B-C**). The *CD68* mRNA transcript level of fluorescence-activated cell-sorted microglia was already elevated two days after ONC and started to decline after day 5 (**Figure 1D**). Microglia mildly reduced their branching when we investigated the morphological branching complexity (**Figure 1E**) and only moderately shifted in the Sholl curve, in contrast to what we have previously seen in the injured eye ^27^. The microglia also did not form any phagocytic cup at the branching tips, typically a sign of a strong microglia response. Finally, we quantified the microglia density in the contralateral eye across the time points since several studies reported an increase ^23,35^. However, the microglia density remained similar across days (**Figure 1F**). Our data suggest that microglia become responsive in the contralateral eye after ONC but do not recapitulate the traditionally phagocytic phenotype observed in the injured eye.

**Figure 1.**
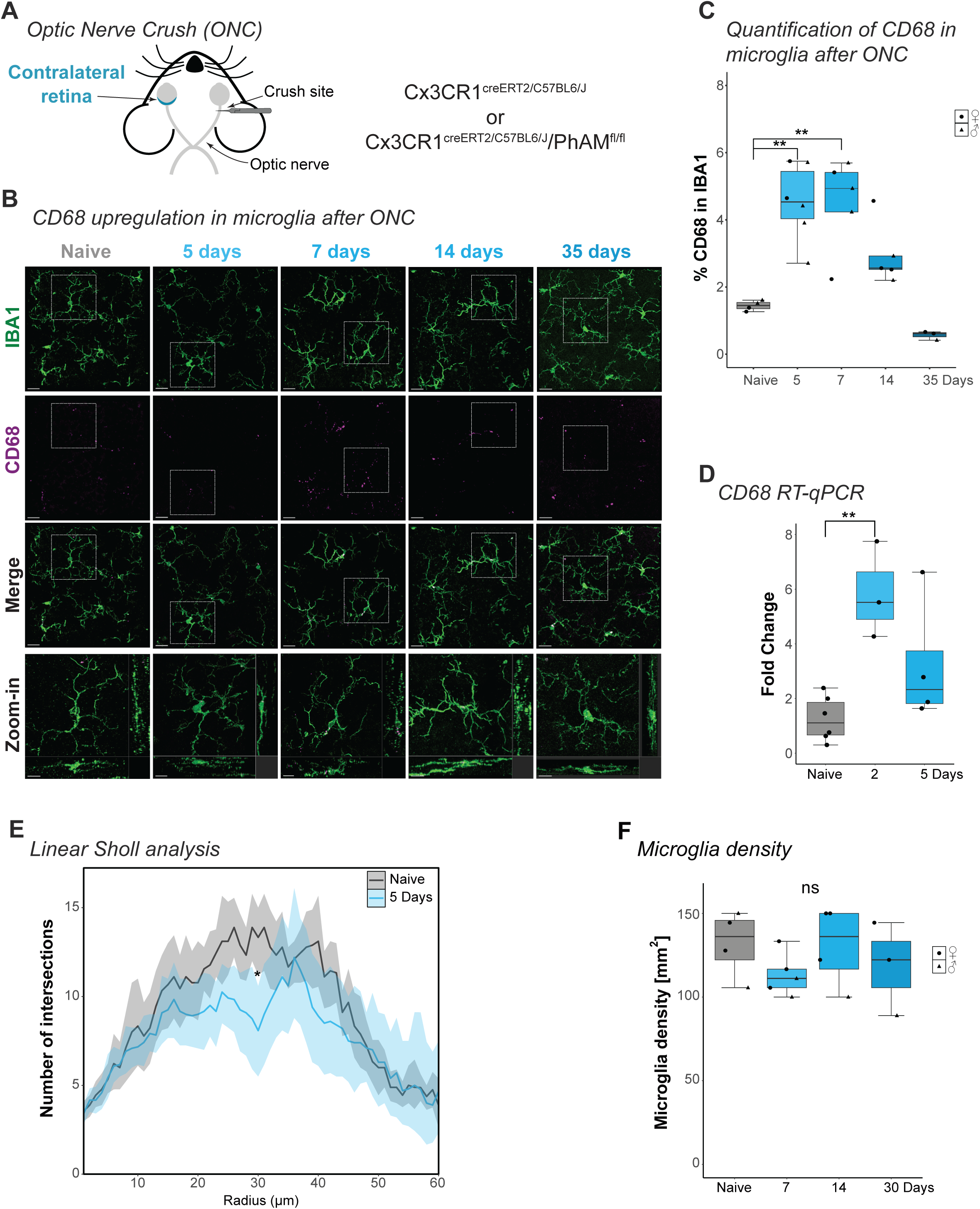
Microglia show a responsive phenotype in the contralateral eye. (A) Schematic of optic nerve crush (ONC). The optic nerve is unilaterally injured using a forceps. (B) Representative wholemount images of the retinal inner plexiform layer (IPL) immunostained for IBA1 (microglia, green) and CD68 (magenta) in naïve or in the contralateral eye 5, 7, 14, or 35 days after ONC. Scalebar: 20 µm, zoom-in with orthogonal projection: 10 µm. (C) Percentage of CD68 volume in IBA1-stained microglia. One-way ANOVA (p= 0.00003, F= 13.18) with shown selected Tukey post hoc: p_naïve *vs*. 5 days_= 0.0012, p_naïve *vs.* 7 days_= 0.0019, p_naïve *vs.* 14 days_=0.213, p_naïve *vs.* 35 days_ = 0.827. (D) Fold change of *CD68* mRNA transcript after ONC compared to naïve. One-way ANOVA (p= 0.008, F= 8.297) with selected Tukey post hoc: p_naïve *vs.* 2 days_= 0.006, p_naïve *vs.* 5 days_= 0.187. (E) Mean number of Sholl interactions per radial distant from the soma (µm) naïve and 5 days after ONC. Linear mixed effect model: p= 0.0392. (F) Microglia density (mm^2^) in the IPL after ONC. One-way ANOVA (p=0.503, F=0.831). For detailed statistical analysis, see **Supplementary Table 1**. **p < 0.01. *p < 0.05. ^ns^p > 0.05, ns= non-significant.

### RBPMS^+^-RGC quantification in the contralateral eye of tamoxifen-induced Cx3cr1^creERT2/C57BL6/J^ mice

Next, we were interested in recapitulating the previously described loss of RGC number ^22–24^. Retinas were collected 35 days after ONC (**Figure 2A**), at which time point the RGCs in the injured retina do not significantly decrease further ^7,40^. The contralateral retina was immunostained for the RGC-marker RBPMS ^41^. To assess the amount of the RGC loss across the retina wholemount in an unbiased approach, we took advantage of the recently developed deep-learning tool RGCode ^32^. The RGCode pipeline automatically detects the contour of the retina wholemount and identifies all RBPMS^+^-cells (**Figure 2B**). As a control, we used naïve Cx3cr1^creERT2^ mice that did not undergo ONC surgery. Surprisingly, the total number of RBPMS^+^-cells was indistinguishable between contralateral and naïve groups (**Figure 2C**). We then normalized the number of RBPMS^+^-cells across the total retinal surface area. This did not change the outcome (**Figure 2D**). Since inflammation might affect the sexes differently, we separated the analysis for female and male mice; however, the results remained the same (**Figure 2E-F**). Our data contradicts the previously described phenotypes ^22–24^.

**Figure 2.**
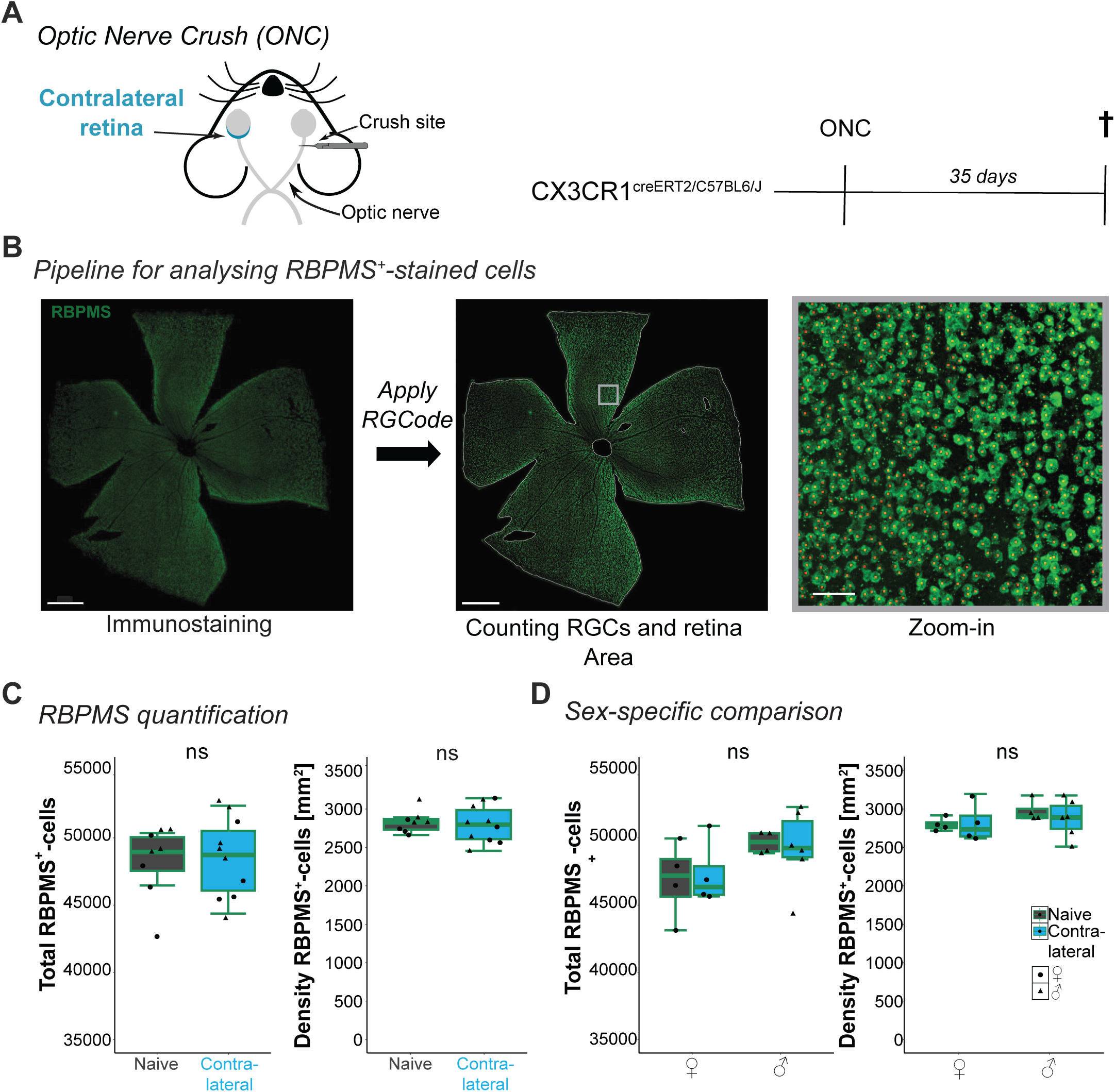
Quantifying RBPMS^+^-RGCs using RGCode in the contralateral eye after ONC. (A) Schematic of optic nerve crush (ONC) and experimental timeline. Tamoxifen injections were performed on CX3CR1^creERT2/C57BL6J^ mice three weeks before ONC surgeries. Retinas were collected 35 days later. (B) Pipeline for analyzing RBPMS^+^-retinal ganglion cells (RGCs) with representative images. Flat-mounted contralateral retinas were immunostained for RBPMS (green) and imaged. RGCode was applied to the entire retina. Zoom-in: RBPMS^+^-labeled RGC overlayed with a detection spot generated by RGCode. Scale bar: 500 µm, zoom-in: 50 µm. (C-D) Box plots of total count and density of RBPMS^+^-cells, including both sexes (C) or separated by sex (D). (C) Student’s t-test for the total count: p=0.8692 (left). Welch test for density: p=0.7348 (right). (D) Total count of RBPMS^+^-cells (left, one-way ANOVA, p=0.301, F=1.342) and density of RBPMS^+^-cells (right, one-way ANOVA, p=0.649, F=0.107). For detailed statistical analysis, see **Supplementary Table 1**. ^ns^p > 0.05, non-significant.

### RBPMS^+^-RGC quantification in the contralateral eye in C57BL6/J mice

To investigate this discrepancy further, we adapted the experiment with wildtype C57BL6/J mouse strain and collected the retinas 45 days after ONC, similar to ^24^ (**Figure 3A**). We quantified the RGC number with RGCode (**Figure 3B**). As before, contralateral retinas from naïve or ONC-injured mice showed similar total RBPMS^+^-RGC counts and RGC density (**Figure 3C**). The RGC count comparisons were unaffected by sex (**Figure 3D**). Since *Cx3cr1*^CreERT2/C57BL6/J^ is a knock-in mouse model, the lack of one Cx3cr1 allele might impact the overall RGC number. Therefore, we compared C57BL6/J and *Cx3cr1*^CreERT2/C57BL6/J^ at corresponding time points. Neither the total RBPMS^+^-cell count nor the RGC density differed between mouse models (**Figure 3E**), excluding the lack of one *Cx3cr1* locus affects contralateral RGC loss.

**Figure 3.**
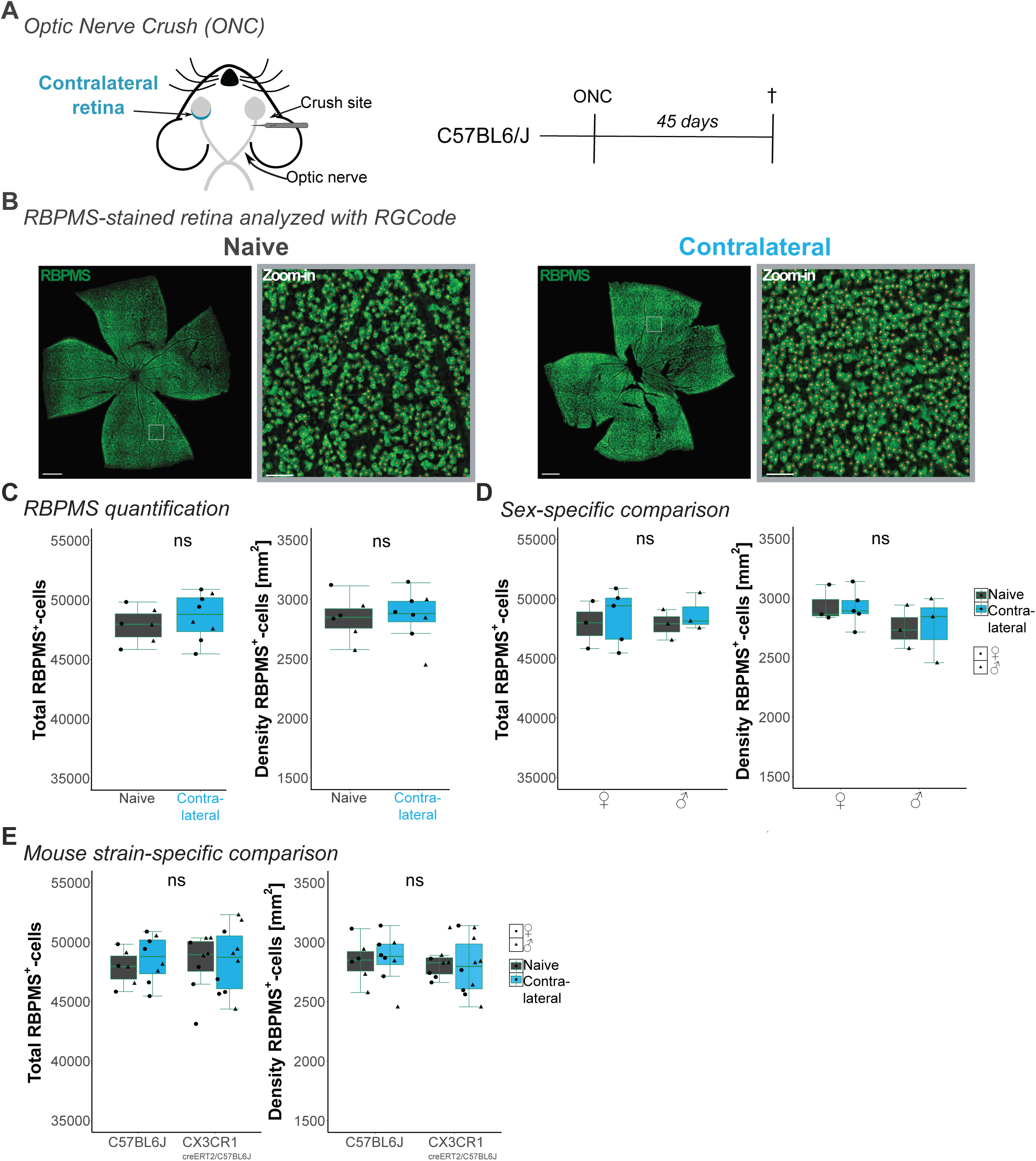
RBPMS^+^-RGCs in the contralateral eye 45 days after ONC. (A) Schematic of optic nerve crush (ONC) and experimental timeline. Retinas of C57BL6/J mice were collected 45 days after ONC. (B) Representative images of flat-mounted retinas immunostained for RBPMS (green) for naïve and contralateral retina. Zoom-in: RBPMS^+^-ganglion cells overlayed with detection spots (red) generated by RGCode. Scale bar: 500 µm, zoom-in: 50 µm. (C-E) Box plots of total count and density of RBPMS^+^-cells, including both sexes (C), separated by sex (D), or mouse strain with CX3CR1^creERT2/C57BL6J^ taken from Figure 2 (E). (C) Student’s t-test for the total count: p=0.4711 (left). Welch test for density: p=0.8733 (right). (D) Box plots comparing influence by sex on total count (left, one-way ANOVA, p=0.917, F=0.165) and density of RBPMS^+^-cells (right, one-way ANOVA, p=0.475, F=0.899) after ONC. (E) Box plots comparing the influence of mouse strain on total count (left, one-way ANOVA, p=0,923 F=0.013) and density of RBPMS^+^-cells (right, one-way ANOVA, p=0.925, F=0.017) in naïve and contralateral retinas. For detailed statistical analysis, see **Supplementary Table 1**. ^ns^p > 0.05, non-significant.

### Development of RGC-Quant to automate Brn3a signature analysis

Besides the mouse strain, we also observed that the studies reporting RGC loss detected RGC with the antibody for the transcription factor Brn3a instead of RBPMS ^24,42,43^. Brn3a is a highly selective RGC marker ^44^, but it has been reported that the expression level acutely decreases when RGCs are under stress ^45,46^. Thus, we hypothesized that detected RGC loss might be an effect of Brn3a expression loss.

Brn3a is localized in the nucleus, while RBPMS labels the cytoplasm (**Figure 4A-B**). The RGCode algorithm is specifically trained to quantify RBPMS^+^-cells and must be retrained to accommodate Brn3a detection in the nucleus ^32^. Moreover, the commonly used Brn3a antibody is raised in mice, which causes some nonspecific background labeling of endogenous IgG in residual blood in the tissue ^47^. This would affect the readout of the deep-learning algorithm.

**Figure 4.**
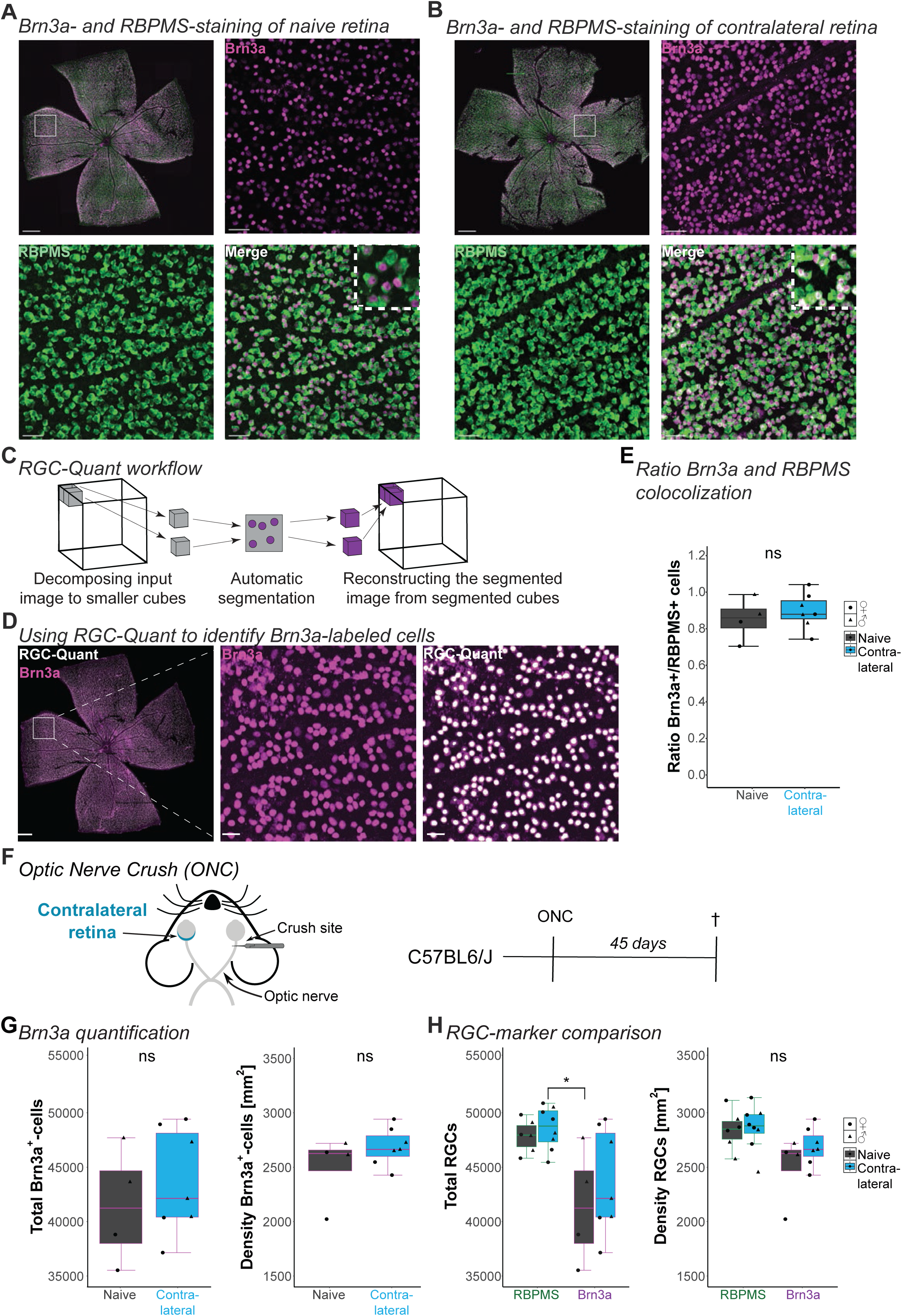
Using RGC-Quant to analyze Brn3a^+^-RGCs in the contralateral eye after ONC. (A-B) Representative images of flat-mounted retinas immunostained for Brn3a (magenta) and RBPMS (green) in naïve (A) and contralateral (B) conditions with zoom-in. Scale bar: 500 µm, zoom-in: 40 and 5 µm. (C) Schematic of the RGC-Quant workflow. 3D images are decomposed into smaller cubes, which are segmented, and the spots are counted. The segmented 3D image is then reconstructed from segmented cubes. (D) Representative image of flat-mounted retina immunostained for BRN3a (magenta) with zoom-in and overlayed with detected spots (white) generated by RGC-Quant. Scale bar: 500 µm, zoom-in: 40 µm. (E) Boxplot showing the ratio of RBPMS-labeled cells co-labeled with Brn3a. Student’s t-test p=0.491. (F) Schematic of optic nerve crush (ONC) and experimental timeline. Retinas of C57BL6/J mice were collected 45 days after ONC. (G-H) Box plots of total count and density of Brn3a^+^-cells (G) and comparison of total count and density for RBPMS and Brn3a (H). (G) Student’s t-test for the total count: p=0.491 (left). Wilcoxon rank test for density: p=0.315 (right). (H) Box plots comparing the influence of antibody on total count (left, Kruskal-Wallis test p=0.0378 with shown selected Conover-Iman post hoc p_Brn3a_naive *vs.* RBPMS_Contralateral_= 0.0239) and density of RGC cells (right, Kruskal-Wallis test p= 0.06501) in naive and contralateral retinas. For detailed statistical analysis, see **Supplementary Table 1**. *p < 0.05. ^ns^p > 0.05, non-significant.

Therefore, we adapted a deep learning model developed in our lab to overcome these challenges. *RGC-Quant* is based on a 3D U-Net architecture ^48^ and was trained using 3D-confocal images of RGC labeled semi-automatically with IMARIS (**Figure S1**). After decomposing the image into small 3D cubes, the model was applied to detect Brn3a^+^-cells automatically in each cube. Once the automatic detection of all the cubes was done, the final segmented image was reconstructed from the small cubes (**Figure 4C**). Combined with post-processing techniques, this method enabled us to detect cells automatically and to compute the number of cells accurately (**Figure 4D**).

### Comparison of Brn3a^+^- and RBPMS^+^-retinal ganglion cells in contralateral retinas

To investigate the possibility that the detected RGC loss might be an effect of Brn3a expression loss, we co-immunolabeled the contralateral retinas of C57BL6/J mice 45 days after ONC with RBPMS and Brn3a (**Figure 4F**). Then, we applied RGCode for the RBPMS^+^-RGC and RGC-Quant for the Brn3a^+^-cells. When we used our algorithm to determine the total number of Brn3a^+^-cells and their density, we saw no significant difference between the naïve and the contralateral condition (**Figure 4G**). Comparing the Brn3a^+^- and RBPMS^+^-labeled retinas in the naïve and contralateral condition, we only found a significant difference between the total number of RGCs in the contralateral retinas stained with RBPMS or Brn3a (**Figure 4H**). This difference between the two markers aligns with the literature that RBPMS labels a larger population of RGC subtypes than Brn3a ^49^. In our dataset, the percentage of Brn3a^+^-cells co-labeled RBPMS^+^-cells is 86 ±12 % for the naïve retinas and 91±10% for the contralateral retinas (**Figure 4F**), aligning with previous findings ^49,50^. Our results suggest no RGC loss after ONC in the contralateral eye, irrespective of the RGC antibody.

## Discussion

This study investigated whether an optic nerve crush (ONC)-induced injury causes RGC loss in the non-injured, contralateral eye. Our study cannot confirm previous data reporting this effect ^22–24^, indicating that methodological differences in quantification might explain this discrepancy.

In recent decades, studies have frequently reported that inflammation is a critical factor in eye injury that does not remain within one eye but also has consequences in the non-injured eye ^25,26^. Macrogliosis and microglia reactivity are common phenotypes when causing injuries such as ONC, optic nerve transection, or increased ocular pressure ^3,16–18^. Even targeting defined retinal cell types with viral approaches like AAV will induce a minor injury because they require either a subretinal or intravitreal delivery route ^51–53^. When we started to investigate this effect, we confirmed a mild inflammatory response based on the microglia phenotype in the contralateral eye, consistent across most studies ^3,16,54^. The only difference that we found was related to the microglia density. Whereas several studies counted more microglia five to seven days after ONC ^17,34,55^, we could not find this effect in the IPL of the contralateral retina, which is in line with other studies ^23,35^. We suspect this discrepancy might be explained by the fact that the microglia density varies across the retina, which might result in a bias when we counted only two quadrants compared to other studies that analyzed the entire retina.

The ambiguity about the RGC loss due to inflammation encouraged us to examine the experimental details of each study closely. We found that the studies differ in the chosen animal model, the investigated time points, and the methods used for RGC labeling. Most studies that report a decrease of RGCs in the contralateral eye have imaged the whole retina in pigmented mice or rat lines. Thus, we focused on two mouse models with a C57BL6/J background and additionally considered sex as a contributor. Sex differences have been shown to affect the progress of neurological diseases with inflammatory components ^56–59^. Our analysis was based on mixed-sex populations, whereas the studies reporting RGC loss in the contralateral eye are primarily performed on males ^22,24^. However, the contralateral eye had no sex-dependent effect.

In contrast to previous research, we could not detect RGC loss in the contralateral retina 45 days after ONC ^24^. This data also indicated no relationship between microglial activation and RGC degeneration in the contralateral eye. Lucas-Ruiz *et al.*,(2019) has also investigated the RGC loss 90 days after ONC, and we were considering looking at this point. However, the authors did not find additional RGC loss compared to 45 days. Furthermore, our data shows that microglia have already lost their reactive phenotype by two weeks, making it unlikely that RGC loss would still be associated with the insult.

Finally, we also explored both Brn3a and RBPMS RGC markers as potential sources for the discrepancy. To reliably visualize RGCs, studies have used antibodies against RBPMS and Brn3a ^46^. The Brn3a/Pou4f1 transcription factor is a nuclear label, delineating the staining well ^45,60^. On the other hand, RBPMS labels more RGC subtypes and, therefore, has become the current gold standard for identifying RGCs ^41,49^. It has been shown that Brn3a is expressed in 80%-90% of the RGCs labeled with RBPMS ^44,50^. Studies suggest that the expression level of Brn3a decreases when RGC are under stress ^45,46^, which could cause the potential difference. Underlining this hypothesis, a study comparing manually counted RBPMS^+^- and Brn3a^+^-RGCs after increased ocular pressure-induced optic nerve damage shows an increased loss of Brn3a^+^-RGCs compared to RBPMS^+^-cells ^61^. Lucas-Ruiz *et al.*,(2019) and Galindo-Romero et al., (2013a) used Brn3a, and, therefore, we compared both antibodies. We confirmed the 80-90% coverage of RBPMS with Brn3a. However, we did not observe the reported RGC loss in the contralateral eye for either of the antibodies.

A primary source of inconsistency is how to quantify RGC loss. Most studies focus on randomly selecting regions of interest in quadrants of the retina and manually or semi-automatic counting the RGC with image analysis software ^20,22,23,35,43^. This analysis is pruned for biases because the RGC density is not uniformly distributed in the murine retina, with higher RGC densities in the central areas close to the optic disc than the periphery and a high-density streak along the nasal-temporal axis ^32,62,63^. Additionally, the ONC-induced RGC loss in the injured eye is diffuse across the entire retina. Therefore, we specifically opted for automated analysis of the whole retina to eliminate user-prone errors induced by manual counting or the positioning of imaging quadrants. RGCode offered a possibility for an unbiased analysis for RBPMS^+^-RGC but would need to be retrained to analyze Brn3a^+^-cells ^32^. Thus, we developed our tool, RGC-Quant, to examine these cells. This deep-learning-based tool can accurately count cells from 3D images, unlike other semi-automatic methods used to analyze Brn3a^+^-RGCs based on 2D single frame images ^24,64^.

An open question that remains is how the ONC-induced microglia phenotype is also detected in the contralateral eye. Several hypotheses for this phenotype have been proposed, such as that the nerve injury compromises the eye’s immune privilege and signals pass through antigen-presenting cells of the immune system ^16,65^. Alternative mediators could be stress signals induced upon ONC and travel into the superior colliculus. There, they cause a responsive microglia phenotype with increased TNF-α levels in both the contra- and ipsilateral side of the ONC ^35,66^. Interestingly, injecting TNF-α into the superior colliculus unilaterally upregulates inflammatory markers and microglial activation in both eyes, suggesting potential retrograde signaling ^66^. Finally, astrocytes could also contribute because they form an extensive coupled network via gap junctions along the optic nerve and the optic chiasm and could connect both eyes ^67,68^. There are still exciting questions about revealing the inflammatory signature communication pathway between the injured and non-injured eye. The insights would have substantial implications in a clinical setting. Nevertheless, our study emphasizes that the contralateral eye should not be used as a control for an ONC experiment. While both eyes have anatomically minimal overlap, they cannot be considered independent.

---

**Figure S1.**
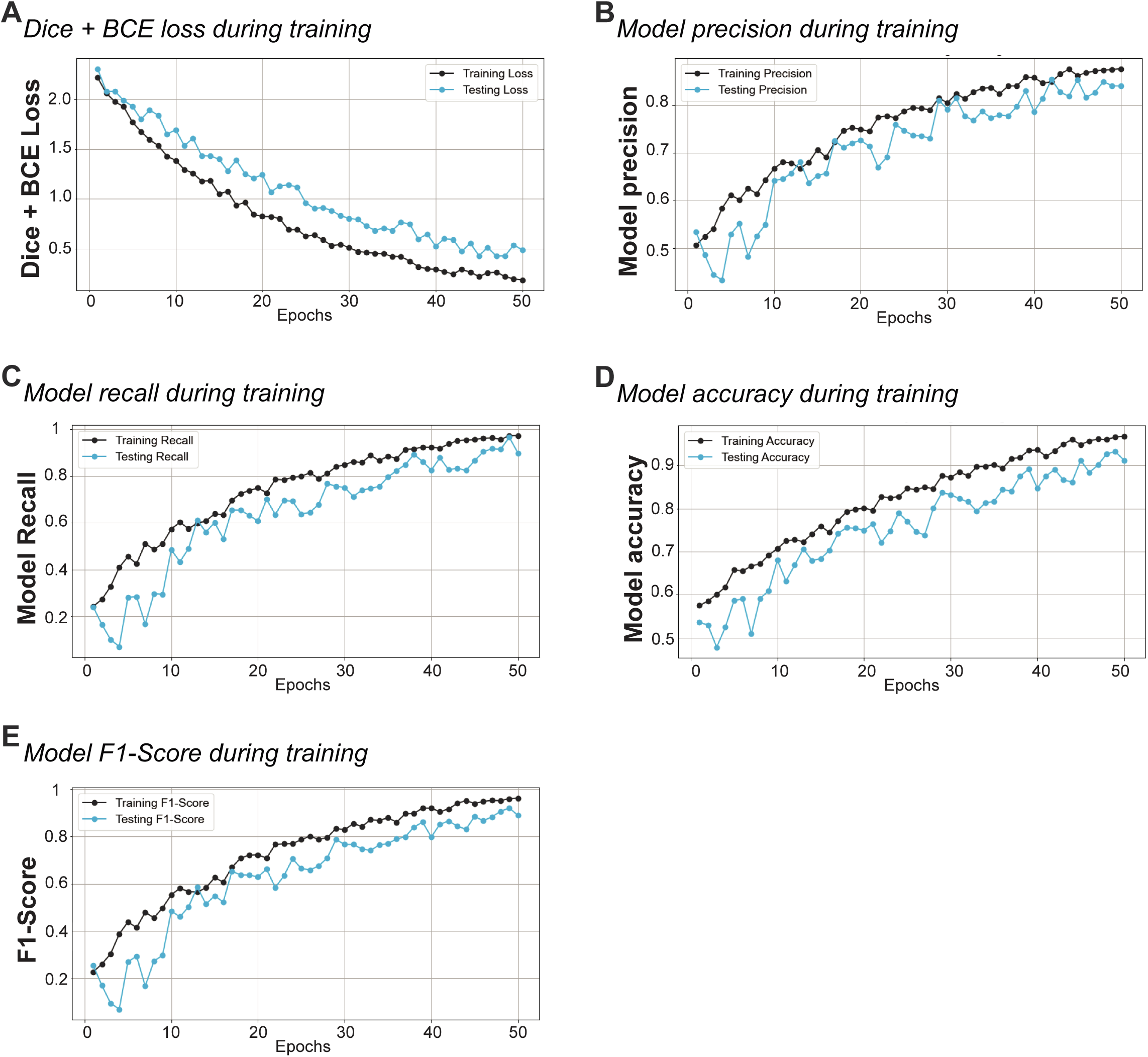
Training of RGC-Quant algorithm. (A) Loss over epochs for training and testing datasets. The loss decreases as the number of epochs, one complete pass of the training dataset through the algorithm, increases, indicating model convergence. BCE: binary cross entropy. (B) Precision over epochs for training and testing datasets. Precision improves as the model learns to minimize false positives. (C) Recall over epochs for training and testing datasets. Recall improves as the model learns to minimize false negatives. (D) Accuracy over epochs for training and testing datasets. Accuracy improves as the model learns to classify more samples correctly. (E) F1-Score over epochs for training and testing datasets. The F1-score, which balances precision and recall, improves as the model continues training.

## Data and code availability

The data reported in this paper are available from the corresponding author upon request. The codes for RGC-Quant are available at https://github.com/siegert-lab/RGC-Quant.

## Acknowledgments

We thank the Scientific Service Units (SSU) of ISTA for the provided resources, specifically the Imaging and Optics Facility (IOF), the Lab Support Facility (LSF), and the Pre-Clinical Facility (PCF) team, specifically Sonja Haslinger, Claudia Gold, and Michael Schunn, for mouse colony management and support. We thank all members of the Siegert group for constant feedback on the project and the manuscript.

## Author contributions

F.S.U., M.E.M, S.S. conceptualized the experimental design. F.S.U. performed experiments, statistical analysis, and created figures. M.E.M. performed ONC, analyzed CD68 content at 5, 14, and 35 days, and performed linear Sholl analysis. G.C. performed FACS-based RT-qPCR experiments and data analysis. A.F. and M.A. developed the RGC-Quant algorithm. F.S.U and S.S. wrote the manuscript and were in charge of project administration. S.S. supervised the project and acquired the funding. All authors commented and discussed the article and approved the final version.

## Notes

### Competing Interest Statement

The authors have declared no competing interest.

